# Spring flowering habit in field pennycress (*Thlaspi arvense*) has arisen multiple independent times

**DOI:** 10.1101/174920

**Authors:** Kevin M. Dorn, Evan B. Johnson, Erin Daniels, Donald L. Wyse, M. David Marks

## Abstract

- Pennycress (*Thlaspi arvense* L.) is currently being developed as a new cold-season oilseed crop. Like many Brassicaceae, pennycress can exhibit either a winter or spring annual phenotype. In Arabidopsis, mutations in negative regulators of flowering, including FLOWERING LOCUS C (FLC) and FRIGIDA can cause the transition to a spring annual habit. The genetics underlying the difference between spring and winter annual pennycress are currently unknown.

- Using whole genome sequencing across wild spring annual pennycress accessions, co-segregation analyses, and comparative genomics approaches, we identify new alleles of TaFLC and explore their geographic distribution.

- We report that loss of function mutations in TaFLC confer the spring annual phenotype in pennycress. We have also identified four natural alleles of TaFLC that confer a spring annual growth habit. The two spring annual FLC alleles present in European accessions were only identified in accessions collected in Montana, USA.

- In pennycress, the spring annual habit has arisen several independent times. Accessions harboring the two European alleles were introduced to North America, likely after the species became a widespread on the continent. These findings provide new information on the natural history of the introduction and spread of the spring annual phenotype in pennycress.

## Introduction

*Thlaspi arvense* L. (field pennycress, pennycress herein) is currently a target for domestication as a new, cold-hardy winter oilseed crop that can fit within the corn/soybean rotation in the Midwestern United States (Dorn *et al.*, 2013; Sedbrook, 2014; Dorn *et al.*, 2015). As a winter annual crop, pennycress can be planted in the late summer to early autumn, either into standing corn or immediately after corn harvest. The pennycress crop establishes a robust vegetative cover prior to winter, providing important ecosystem services such as limiting soil erosion and nutrient runoff. In the spring, pennycress will flower and set seed, yielding upwards of 1,600 pounds per acre of oil-rich seed in time for planting a crop of soybean (Phippen & Phippen, 2012; Sedbrook, 2014). Pennycress seed is high in oils that can be readily converted to biodiesel or jet fuel, with the remaining high-protein seed meal having potential as animal feed or feedstock for industrial uses (Moser *et al.*, 2009; Selling *et al.*, 2011; Evangelista *et al.*, 2012; Fan *et al.*, 2013; Hojilla-Evangelista *et al.*, 2013).

Pennycress is a member of the Thlaspideae, a tribe of the Brassicaceae (Beilstein *et al.*, 2008). The Brassicaceae family is divided into three lineages. Pennycress resides in lineage II, along with the Brassica genus, including the economically important oilseed crops *Brassica rapa* and *Brassica napus*, but not the model species *Arabidopsis thaliana* nor *Capsella rubella*, which are members of lineage I (Franzke *et al.*, 2011). Pennycress is native to Eurasia and is considered naturalized to most temperate to subarctic regions throughout the northern hemisphere, including all of the United States except for Hawaii and Alabama, all provinces of Canada, as well as being considered naturalized in the southern hemisphere, including Australia, New Zealand, and Argentina (Warwick, 2002). Pennycress exhibits numerous traits that make it an attractive winter rotation crop (Sedbrook, 2014). Of particular importance in the upper Midwest is the existence of pennycress lines that complete their life cycle rapidly enough to fit within the corn/soybean rotation. Central to this rapid spring development of pennycress are its underlying growth habits. In wild populations, there are both winter and spring annual pennycress (Best & McIntyre, 1972; McIntyre & Best, 1978), similar to many other Brassicaceae species such as *Arabidopsis thaliana* (Stinchcombe *et al.*, 2004; Shindo *et al.*, 2005), *Brassica rapa* (Wu *et al.*, 2012), *Camelina sativa* (Crowley, 1999), and *Brassica napus* (Tadege *et al.*, 2001).

Throughout the decades of research on basic developmental questions in Arabidopsis, and the expanding translational research in other Brassica crops, the underlying molecular mechanisms controlling flowering time have been identified in many of these species (Simpson & Dean, 2002; Amasino, 2005; Jung & Muller, 2009; Kim *et al.*, 2009). In wild accessions of Arabidopsis, only a handful of mutations are responsible for variation of flowering time and vernalization requirement (Burn *et al.*, 1993; Clarke & Dean, 1994; Johanson *et al.*, 2000). Most notably, allelic variation in two key negative regulators, FLOWERING LOCUS C (FLC) and/or FRIGIDA (FRI), underlies the key difference between spring and winter annual plants (Johanson *et al.*, 2000; Michaels & Amasino, 2001; Michaels *et al.*, 2003; Michaels *et al.*, 2004; Shindo *et al.*, 2005; Jiang *et al.*, 2009). FLC encodes a MADS box transcription factor (Michaels & Amasino, 1999), and inhibits flowering prior to vernalization by repressing the expression of FLOWERING LOCUS T (FT) (Searle *et al.*, 2006). In Arabidopsis, *flc* null mutations eliminate the vernalization requirement and impart the spring annual, rapid-flowering phenotype (Michaels & Amasino, 1999). Allelic variation within FRI also impacts flowering time and the vernalization requirement. Similar to FLC, Arabidopsis accessions harboring loss of function mutations in FRI flower rapidly without vernalization (Johanson *et al.*, 2000; Shindo *et al.*, 2005).

The expression of FLC is positively regulated by FRI (Michaels & Amasino, 1999), thus, mutations in either of these vernalization-responsive negative regulators can lead to a loss of vernalization requirement and a spring annual growth habit (Michaels & Amasino, 1999; Michaels & Amasino, 2001). The vernalization signal provided by the cold of winter removes the repression on the transition to flowering through the epigenetic silencing of FLC. Specifically, vernalization increases histone 3 K27 trimethylation (H3K27me3) at FLC chromatin, reducing transcriptional activity (Sung *et al.*, 2006; Finnegan & Dennis, 2007; Greb *et al.*, 2007; Coustham *et al.*, 2012; Yang *et al.*, 2014). The vernalization-induced silencing of FLC releases the repression on the transition to flowering, which permits the transition to reproductive growth (Searle *et al.*, 2006). Additionally, the roles of the COOLAIR long noncoding antisense transcripts transcribed from the AtFLC locus have been revealed as a key regulatory component in the cold-induced regulation of FLC (Swiezewski *et al.*, 2009; Csorba *et al.*, 2014; Marquardt *et al.*, 2014).

While this extensive understanding of the molecular genetic pathways controlling flowering time and vernalization in Arabidopsis has informed similar studies in Brassica relatives, little is known about the underlying mechanisms controlling flowering time variation in pennycress. Different accessions of pennycress have been reported to either act as early flowering or late flowering, with late flowering accessions growing as rosettes for as long as 150 days prior to flowering (Best & McIntyre, 1972). It was later found that vernalization increased the rate of flowering in the late flowering accessions (Best & Mc Intyre, 1976). Analyses of F2 progeny between the late and early flowering accessions determined the early flowering phenotype (spring annuals lacking the vernalization requirement) was determined by a single gene, with the late flowering allele being completely dominant (McIntyre & Best, 1978).

In this report we address the questions concerning the molecular basis for the spring flowering habit in pennycress, whether or not the spring habit has arisen only once or multiple times in nature, and we begin to analyze the geographic distribution of pennycress flowering time variants. These studies were aided by genomic resources that have been developed for pennycress, including a transcriptome and draft genome (Dorn *et al.*, 2013; Dorn *et al.*, 2015) These resources capture the pennycress gene space and greatly aided in the identification of alterations that lead to spring flowering. We show that key mutations in the FLC gene are responsible for spring flowering in many natural pennycress accessions, including one allele of FLC that confers the spring annual growth habit that spread from Europe to North America.

## Materials and Methods

### 1.) *Thlaspi arvense* accessions and F2 population

The spring annual *Thlaspi arvense* line MN108SA was derived from a wild Minnesota population containing both winter and spring annual plants. Five generations of single seed descent was performed on a spring annual plant from this collection and sequenced. A single MN111 plant was also carried through two generations of single seed descent and sequenced.

Additional pennycress accessions described here with the ‘PI’ or ‘Ames’ prefix were obtained from USDA-GRIN. All pennycress accessions with the ‘MN’ prefix were collected by Dr. Donald Wyse (University of Minnesota). The spring annual line Spring32 was obtained from Dr. Win Phippen at Western Illinois University under a Materials Transfer Agreement.

Plants were germinated on moist Berger BM2 germination mix (Berger, Inc., www.berger.ca), stratified at 4**°** C for 7 days, and grown in climate-controlled growth chambers at the University of Minnesota (21**°** C, 16 hour/8 hour day/night cycles at 100 micromoles/m^2^/s PAR). The MN111 plant sequenced in this analysis was vernalized at six weeks post-germination at 4**°** C for 30 days in the dark. After vernalization, this plant was returned to the growth chamber conditions described above. The spring annual accessions were not vernalized as they flowered immediately.

### 2.) DNA isolation and Illumina genomic DNA sequencing

DNA was isolated from a single MN108SA and a single MN111 plant using the Omega Mag-Bind Plant DNA kit (Omega Bio-Tek, www.omegabiotek.com) according to the manufacturer’s recommended protocol. These DNA samples were submitted to the University of Minnesota Genomics Center for sequencing on the Illumina HiSeq 2000 platform (Illumina Inc, www.illumina.com). Sequencing libraries were prepared using the Illumina TruSeq Nano DNA Sample Prep kit with an average library insert size of 460 base pairs. The Illumina Universal Adaptor and an Indexed Adaptor (MN111 – Illumina Indexed Adaptor #12 – barcode = CTTGTA, MN108SA – Illumina Indexed Adaptor #11 – barcode = GGCTAC) were used to create the sequencing libraries. Each library was sequenced on a full lane of Illumina HiSeq 2000 (100 base pair, paired-end).

DNA from the six selected European pennycress accessions was isolated using the Qiagen Plant DNeasy Mini kit (www.qiagen.com) following the manufacturer’s recommended protocol. Illumina sequencing libraries were created for each European accession using the Illumina TruSeq Nano kit with the 350 bp insert protocol. These six libraries were sequenced across 1.5 lanes on the Illumina HiSeq 2500 platform (125 base pair, paired-end) with version 4 chemistry.

All raw sequencing datasets have been deposited in the NCBI Short Read Archive under BioProject PRJNA237017 / SRA Accession SRP036068 (Table S1).

### 3.) Sequencing data quality control, read mapping

Illumina sequencing read quality and contamination were examined using FASTQC (http://www.bioinformatics.babraham.ac.uk/projects/fastqc/). FASTQ files were filtered and trimmed to remove low quality reads and sequencing adaptors using BBDuk (https://sourceforge.net/projects/bbmap/) using the following parameters: ftl=10 minlen=50 qtrim=rl trimq=10 ktrim=r k=25 mink=11 hdist=1 ref=/bbmap/resources/adapters.fa tpe tbo. Sequencing reads from accessions MN111 and MN108SA had the additional BBDuk flag of ftr=95. The resulting high quality reads from each accession were mapped to the v1 pennycress genome using Bowtie 2 and visualized in CLC Genomics Workbench.

### 4.) Analysis of FLC mutations in MN111 x MN108SA F2 population, global spring varieties, and EMS mutant

DNA was isolated from F2 progeny grown in the conditions described above using the Omega Bio-Tek Plant MagBind 96 kit according to the manufacturers recommended protocol. DNA oligos were designed to amplify the 5’ end of the pennycress FLC gene for the *flc-A* allele, approximately 100 base pairs upstream of the transcriptional start site (TaFLC_1_Forw: 5’ – CCGAGGAAGAAAAAGTAGATAGAGACA – 3’, TaFLC_1_Rev: 5’ – GAAGCTTAAAGGGGGAAAAAGGAA – 3’, Table S2). This amplicon was also able to identify the *flc-D* allele in MN133 and PI633414 and the *flc-*α allele from the EMS-mutagenized population as well. Polymerase Chain Reaction (PCR) was used to amplify this fragment, producing an approximately 450 bp amplicon. New England Biolabs Q5 High-Fidelity PCR Kit with 2x Master Mix was used, with the following thermal cycler conditions: 1.) 98**°** C for 30 seconds, 2.) 98**°** C for 10 seconds, 3.) 57**°** C for 20 seconds, 4.) 72**°** C for 20 seconds, 5.) Go to step #2 34 times, 6.) 72**°** C for 2 minutes, 7.) 4**°** C hold. Reactions were visualized to confirm amplification of a single band using gel electrophoresis. PCR products were submitted to Beckman Coulter Genomics for PCR product purification and single pass Sanger sequencing. Amplicons were sequenced in both directions using the forward and reverse primers listed above. Sanger sequencing reads were analyzed in CLC Genomics Workbench and aligned against the pennycress MN106 reference genome at the FLC locus to identify sequence variants.

### 5.) Identification of deletion alleles using PCR

A diagnostic PCR test was designed to test for the *flc-B* allele. A primer set (TaFLC_2_Forw – 5’ – ATAGTGTGCATCAACTGGTC – 3’, and TaFLC_2_Rev – 5’ – CGAACCATAGTTCAGAGCTT – 3’, Table S2) were designed to amplify a single amplicon overlapping the 456 bp deletion (Fig. S1). In the absence of the *flc-B* allele, a 2,088 bp fragment is produced, whereas a 1,632 bp fragment is produced in plants containing the *flc-B* allele (Fig. 2B). New England Biolabs Q5 High-Fidelity PCR Kit with 2x Master Mix was used, with the following thermal cycler conditions: 1.) 98**°** C for 30 seconds, 2.) 98**°** C for 5 seconds, 3.) 63**°** C for 15 seconds, 4.) 72**°** C for 60 seconds, 5.) Go to step #2 34 times, 6.) 72**°** C for 2 minutes, 7.) 4**°** C hold.

A diagnostic PCR test was developed to test for the presence or absence of the *flc-C* allele (Fig. 2C, Fig. S2). A primer set (TaFLC_4_Forw: 5’ – GCGACGGTGAATATGGAGTTGG – 3’, and TaFLC_3_Rev – 5’ – GCTAATTTTTCAGCAAATCTCCCG – 3’, Table S2) was designed to amplify a 6,598 bp amplicon overlapping the 4,724 bp deletion. In plants containing the *flc-C* allele, this region produces a 1,874 bp amplicon, confirming the presence of this deletion. New England Biolabs Q5 High-Fidelity PCR Kit with 2x Master Mix was used, with the following thermal cycler conditions: 1.) 98**°** C for 30 seconds, 2.) 98**°** C for 5 seconds, 3.) 67**°** C for 15 seconds, 4.) 72**°** C for 40 seconds, 5.) Go to step #2 34 times, 6.) 72**°** C for 2 minutes, 7.) 4**°** C hold. This amplicon was purified using the Qiagen QiaQuick PCR Purification Kit (www.qiagen.com) from accessions MN135, Ames22461, and PI650285 and subjected to Sanger sequencing to confirm the exact deletion size and coordinates. Identical 4,724 bp deletions were identified in these three accessions.

A secondary amplicon was used to confirm the absence of this deletion, with the forward primer (TaFLC_1_Forw: 5’ – CCGAGGAAGAAAAAGTAGATAGAGACA – 3’, Table S2) lying in the middle of the deleted region (Fig. S2). Successful amplification using this secondary primer set (TaFLC_1_Forw and TaFLC_3_Rev – 5’ – GCTAATTTTTCAGCAAATCTCCCG – 3’, Table S2) produces a 1,397 bp fragment and confirms the absence of this deleted region. New England Biolabs Q5 High-Fidelity PCR Kit with 2x Master Mix was used, with the following thermal cycler conditions: 1.) 98**°** C for 30 seconds, 2.) 98**°** C for 5 seconds, 3.) 64**°** C for 15 seconds, 4.) 72**°** C for 40 seconds, 5.) Go to step #2 34 times, 6.) 72**°** C for 2 minutes, 7.) 4**°** C hold.

## Results

### Flowering time phenotypes of winter and spring annual pennycress

While identifying germplasm to be used for the pennycress breeding program, wild isolates of pennycress have been collected from various locations across North America (Table S3). A spring flowering pennycress variant, named MN108SA, was isolated from a mixed population of both spring and winter annual individuals collected from Roseau, Minnesota. Single seed descent for 5 generations showed that the trait was stable and that vernalization was not required for floral induction. MN108SA was crossed to the vernalization-requiring accession MN111. A comparison of MN108SA and MN111 seedlings is shown in Fig. 1A. MN108SA shows internode elongation soon after the formation of the first true leaves (Fig. 1A insert), whereas MN111 forms a rosette. As previously reported by McIntyre and Best, 1978, the F1 derived from the cross between MN111 and MN108SA exhibited the dominant winter flowering habit, segregating in the F2 population at 38:12 winter annual:spring annual, suggesting that a single dominant gene is responsible for maintaining the winter annual phenotype.

**Fig. 1:**
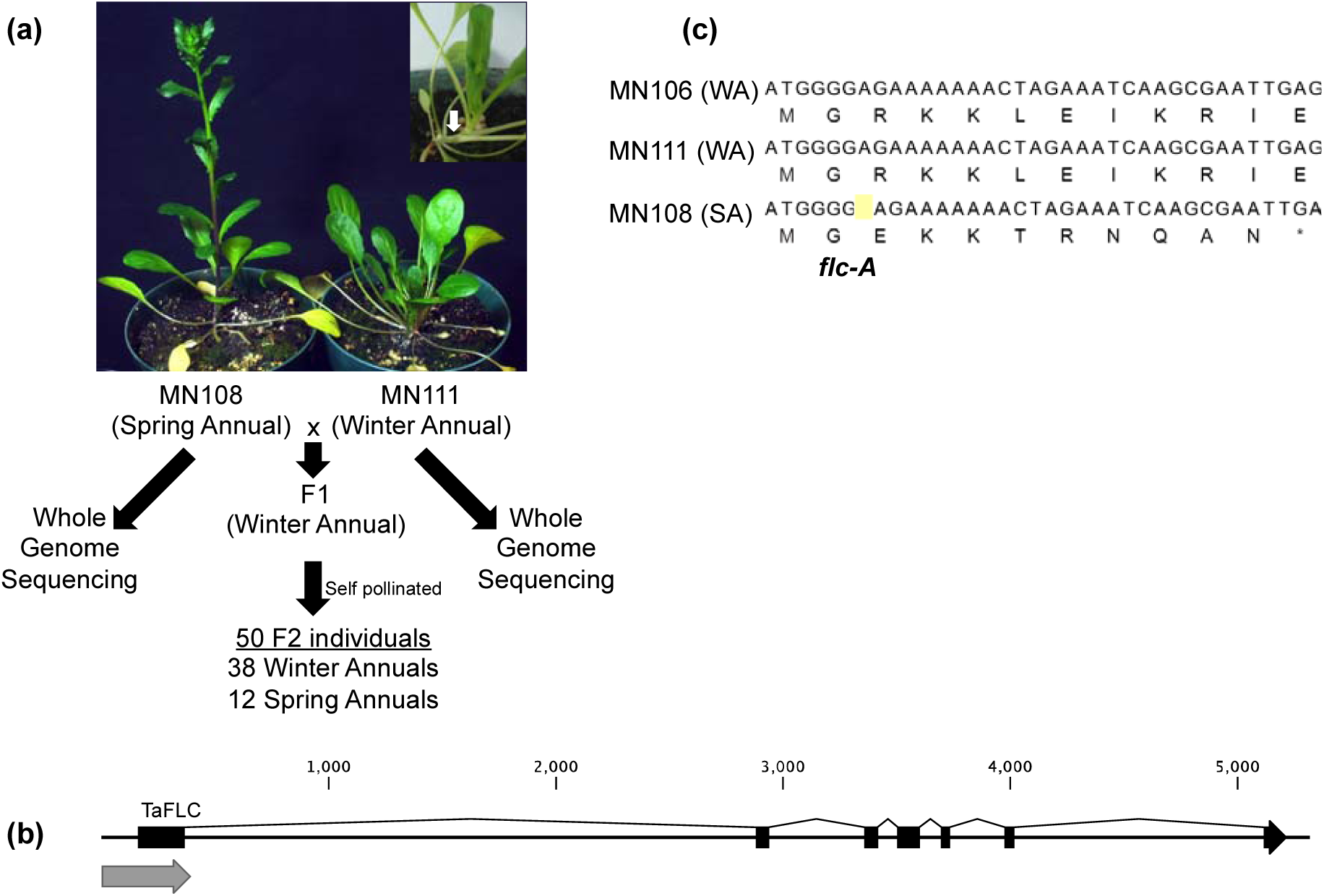
Segregation analysis confirms *flc-A* allele confers spring annual phenotype. (a) Co-grown MN108SA and MN111 pennycress plants. Insert shows immediate internode elongation in the MN108SA spring annual plant. An F2 population was derived from a cross between MN108SA x MN111 and the segregation ratio of spring to winter annual phenotypes was determined. Whole genome sequencing of the parent plants from each background was also conducted. (b) Gene model of pennycress FLOWERING LOCUS C (FLC). Gray arrow below the first exon indicates the position of the amplicon used to genotype the SNP shown in panel C. (c) Nucleotide and predicted peptide sequence of the 5’ end of the pennycress FLC gene in winter annual MN106, winter annual MN111, and the spring annual accession MN108SA. The *flc-A* mutation in MN108SA is highlighted in yellow.

Whole genome sequencing of the MN111 and MN108SA parent plants was used to identify key genomic differences underlying the difference in habit between winter annual and spring annual plants. DNA from each parent was sequenced on a full lane of the Illumina HiSeq 2000 platform (100 base pair, paired end). Over 334M sequencing reads were generated per parent (Table S1). A total of 29B high quality (post quality control) were generated per parent, representing >50X coverage of the predicted genome size of 539Mb ^1^. Trimmed and filtered reads were mapped to the winter annual MN106 draft genome sequence ^2^.

### Whole genome sequencing identifies the pennycress *flc-A* allele

Read mappings at the pennycress FRI and FLC loci for MN106, MN111, and MN108SA were examined to identify potential mutations underlying the winter to spring annual transition. Comparison of the pennycress FRI homolog (Ta1.0_26225 on scaffold 1344 of the v1.0 pennycress genome) revealed a single SNP between these three lines, however, this SNP was shared by both the MN111 and MN108SA individual, likely indicating a non-causative effect on the spring annual growth habit (Fig. S3). This SNP was found to cause an amino acid change (Threonine to Serine) at position 553 of the TaFRI predicted peptide (position 2,085 of the gene model-c.2,085A>T).

Within the MN106 draft genome, a single copy of FLC with a high degree of similarity to that of Arabidopsis was identified (Fig. 1B, Fig. S4, Fig. S5). We identified a single guanine base insertion in the 7^th^ position of the FLC coding sequence (*flc-A*) of the spring annual MN108SA accession that was absent in both winter annual accessions. This frameshift mutation caused a predicted TGA stop codon in the 12^th^ codon position (Fig. 1C).

To obtain genetic evidence that this *flc-A allele* was the causal variant responsible for the spring flowering habit in MN108SA, an F2 population from the cross between MN111 and MN108SA was examined. A single F1 individual was self pollinated, and 50 F2 progeny were planted for further analysis. Of these F2 individuals, 38 exhibited a winter annual phenotype and 12 exhibited a spring annual phenotype (Fig. 1A).

PCR primers were developed that amplify a 394 bp amplicon that covers the location of the *flc-A* insertion. Amplicons from all 12 spring annual F2 progeny were sequenced and confirmed homozygous for this *flc-A* allele. Amplicons from a random selection of 14 winter annual F2 progeny were also sequenced. Of the winter annual F2 progeny, 5 were homozygous for the wild type FLC gene while 9 were heterozygous. A chi square analysis (χ2=0.026667, p-value = 0.8703) indicates that the data fits a 1:2:1 segregation pattern that is predicted for a single recessive mutation in this F2 population.

### Identifying a novel EMS-induced allele of FLC in pennycress

As a component of ongoing work to domesticate pennycress, an EMS-mutagenized MN106 population has been generated (Sedbrook, 2014). This winter flowering population has been screened to identify early flowering variants. In the course of this screen, a new spring flowering time mutant was identified (Fig. S6). This line did not require vernalization and flowered immediately upon germination. Whole genome sequencing of this mutant revealed a novel mutation in the first exon of FLC (*c.52C>T*), resulting in an immediate stop codon and truncated predicted protein (Fig. S6), which was designated *flc-*α. The above segregation data along with the finding of two new independent spring flowering mutants with FLC mutations confirms the causative role of FLC mutations in the spring flowering habit in pennycress.

### Global distribution of three additional spring annual alleles of FLC

With the genetic confirmation that the *flc-A* allele was indeed the causal variant responsible for the spring annual growth habit in MN108SA, we next examined the distribution of this allele in spring annual accessions from around the world. We screened a total of 35 spring annual accessions, which were obtained either from the University of Minnesota collections or GRIN. Of these, 23 were confirmed as pure spring annual accessions in both field and growth chamber experiments, with no winter annual segregation nor contaminating winter annual seed (Table S3). Surprisingly, we only identified the *flc-A* allele in three other accessions, all from North America (MN131 from Montana, USA and Ames31489 and Ames31491 from Canada).

As there were a limited number of representative spring annual accessions in our collection containing the *flc-A* allele, we utilized a PCR-based approach to determine whether there were additional mutations in FLC underlying the spring annual growth habit. As we had already amplified and sequenced the first exon of FLC in all of our spring annual accessions, we moved next to the second exon. Interestingly, we were unable to amplify the second exon in several spring annual accessions. Thus, we expanded our PCR experiments into the flanking introns around exon 2, which lead to the discovery of a second allele of FLC in spring annual lines. This mutation was characterized by a 456 bp deletion that encompasses the entire second exon of FLC, which was named *flc-B*. This allele was identified in 9 spring annual accessions distributed across the United States and Canada, along with the Spring32 accession from the Western Illinois University breeding program (Fig. 2B and 2E, Table S3) (Sedbrook, 2014).

**Fig. 2:**
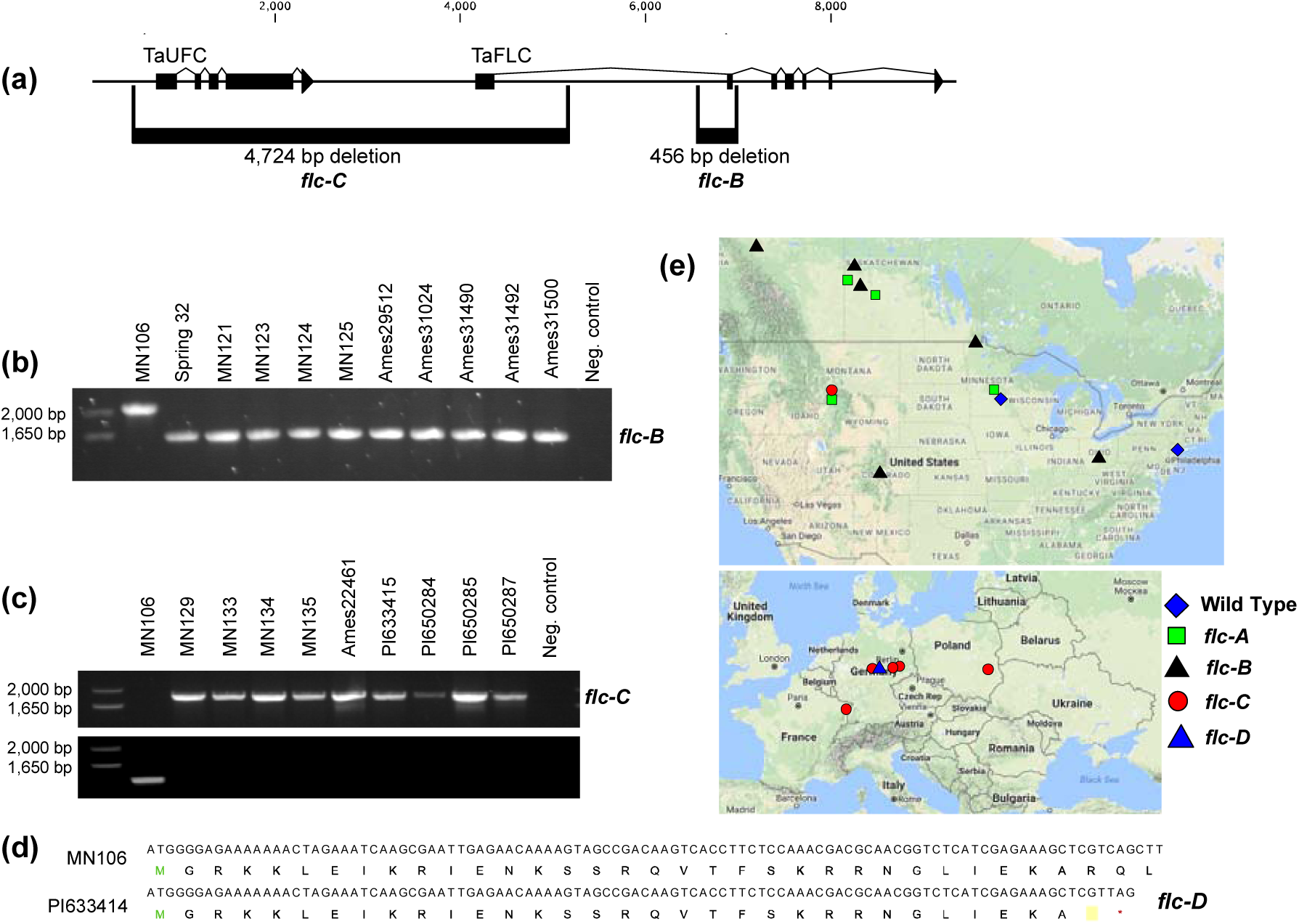
Identification of *flc-B* and *flc-C* alleles of FLC. (a) Expanded gene model of pennycress FLC region, including the upstream TaUFC gene model. Locations of the *flc-B* and *flc-C* alleles are shown below the gene model. (b) Agarose gel of the diagnostic polymerase chain reaction (PCR) results testing for the *flcB* allele across the shown pennycress accessions. The wild type MN106 allele generates a 2,088 bp amplicon, where as the *flc-B* allele produces a 1,632 bp amplicon (Fig S1). (c) Agarose gel of the diagnostic polymerase chain reaction (PCR) results testing for the *flcC* allele across the shown pennycress accessions. Amplification in the top row confirms the presence of the deletion, whereas amplification in the second row confirms absence of the deletion. The sequences of primers used are shown in Table S2, and primer locations and amplicon strategy is shown in Fig. S2. Results for accession PI650286 (Groitzsch, Saxony, Germany) are not shown, but was confirmed to have the *flc-C* allele. (d) Nucleotide and predicted peptide sequence of FLC in MN106 and PI633414 (Wachstedt, Germany), with the *flc-D* allele highlighted in yellow. (e) Global distribution of the four spring annual alleles of FLC in pennycress accessions tested in this study (listed in Table S3).

Having not yet identified a candidate spring annual FLC allele in any accession from Europe, we again employed whole genome sequencing of the 5 publicly available spring annual pennycress accessions (collected in Germany, Poland, and France). Using this approach, we were able to quickly identify a third allele, characterized by a 4,724 bp deletion (*flc-C*), which removes the first intron of FLC along with the upstream gene (gene model Ta000918), a homolog of the Arabidopsis gene UPSTREAM OF FLC (UFC, AT5G10150). In Arabidopsis, UFC and FLC are co-regulated, however, UFC is not believed to influence the vernalization response (Finnegan *et al.*, 2004; Andersson *et al.*, 2008). A diagnostic amplicon experiment was designed to screen the remaining unknown spring annual accessions for the *flc-C* allele. This allele was identified in four accessions collected in Howard Springs, Montana, USA, along with three accessions from Germany, as well as accessions from Poland and France (Fig. 2C, 2E, Supplemental Table 3).

A fourth allele, *flc-D*, was identified through whole genome sequencing of the spring annual accession PI633414 (Germany). This allele is characterized by a single base pair change at the 100^th^ position of the CDS and caused a precocious TAG stop codon in the predicted protein sequence (Fig. 2D). As the position of this mutation was covered by the 394 bp amplicon used to identify the *flc-A* allele, we re-examined the Sanger reads from all of the accessions sequenced in our search for accessions containing the *flc-A* allele. We did not identify any additional accessions containing the *flc-D* allele.

## Discussion

### Use of NGS to quickly identify mutations in FLC

At the onset of this experiment, the predicted costs associated with PCR-based cloning and sequencing of all candidate genes was assessed. This analysis indicated that it is now less expensive to use the whole genome sequencing (WGS) approach to first identify a candidate locus in the parent accessions, and then proceed to test F2 progeny via PCR and Sanger sequencing. Here we report the discovery of four natural alleles of FLC that confer the spring annual phenotype in pennycress. The added value with the WGS approach here is that we now also have genome-wide markers for the two parents of the F2 mapping population (MN111 and MN108SA), which is being used to develop a linkage map via Genotype by Sequencing (Elshire et al., 2011; Poland & Rife, 2012). Additionally, as the network controlling flowering time is known to be extremely complex, the gene variants in each of these parents that may have a minor effect on vernalization and flowering provide a wealth of untapped information for later investigation.

### Global distribution of FLC alleles and the introduction to North America

Pennycress is native to Europe and Asia. Thought to have been introduced to North America as early as the establishment of Fort Detroit (Michigan, USA) in 1701, pennycress became widely distributed throughout Canada and the United States by the early 1900s (Best & McIntyre, 1975). Despite the limited number (35) of pure spring annual accessions tested in this study, we can begin examining the origins of each of the spring annual alleles of FLC identified here.

As the species originated in Eurasia and only the *flc-B and flc-D* alleles were identified in the seven spring annual European accessions tested here, we hypothesize that these alleles originated prior to introduction to North America (Fig. 2E). As the accessions collected near Howard Spring, Montana, USA (MN129, MN133, MN134, MN135) contained one of these ‘European’ alleles of FLC (*flc-C*), this suggests that this allele may have been introduced into North America a limited number of times, potentially from plants near Saxony or Thuringia, Germany (Fig. 2E). However, it is possible that the *flc-D* allele is present in populations in North America, but was not identified in the accessions represented in this experiment.

Similarly, as we did not identify European accessions containing the *flc-A* or *flc-B* alleles, we predict that these alleles may have originated after the introduction of pennycress to North America and this allele is unique to North American accessions. It is possible that there are in fact European pennycress populations that contain the *flc-A* allele, but with the limited European germplasm availability, were not found in this study. In this scenario, it is feasible that a mixed seed lot containing *flc-A* and *flc-B* alleles were introduced to Montana in a single event, and plants with the *flc-A* and *flc-B* alleles spread significantly more compared to those with the *flc-D* allele.

The accessions collected near Howard Spring, Montana (MN129, MN131, MN133, MN134, MN135) are an interesting case study of these hypotheses, especially considering the findings of the *flc-A* allele in MN131, the *flc-B* allele in MN129, MN133, MN134, and MN135, and a lack of identification of European accessions with the *flc-A* allele. The above scenarios could be investigated further in these accessions collected from Montana using genome-wide markers from either genotype-by-sequencing or WGS. As we have already generated WGS data for five of the European accessions in this study, sequencing of these accessions from Montana and a comparison of shared polymorphisms may help in determining the relatedness between the North American and European lines, and may point to the exact origins of these alleles of FLC in pennycress. Again, with the limited number of pure spring annual accessions available, future studies should also focus on the collection of a more comprehensive set of globally distributed lines, including both spring and winter annuals.

### Towards an understanding of flowering time control in pennycress

Our previous efforts to develop genomic resources for the domestication of pennycress identified likely homologs controlling flowering time and the vernalization response via DNA and RNA sequence homology (Dorn *et al.*, 2013; Dorn *et al.*, 2015). Here we present the first sequence-supported genetic information on the underlying mechanisms controlling flowering time in pennycress. While the four natural alleles of FLC reported here are sufficient to confer the spring annual phenotype, there is likely a host of interacting genes and epigenetic factors also essential for the rapid flowering seen in the spring annual accessions investigated here. Of particular interest is FRIGIDA and members of the ‘FRIGIDA Complex’ (FRI-C), including FRIGIDA LIKE 1 (FRL1), FRIGIDA-ESSENTIAL 1 (FES1), SUPPRESSOR OF FRIGIDA4 (SUF4), and FLC EXPRESSOR (FLX) (Choi *et al.*, 2011). The FRIGIDA Complex acts as a transcriptional activation complex on FLC expression through a diverse range of functions of each complex member (Choi *et al.*, 2011). While no loss of function FRIGIDA alleles were identified in the accessions examined here, it remains possible that an FLC-independent path to the spring annual phenotype exists in light of the limited availability of globally distributed germplasm. The interaction between the vernalization response, photoperiodic, and autonomous pathways are also of great interest, as allelic variation and unique combinations of alleles from each pathway can contribute to quantitative variation in flowering time and lead towards the goal of developing an elite winter annual pennycress variety that flowers rapidly in the spring to fit within the short growing season of northern climates.

## Acknowledgements

The authors would like to thank Ryan Emenecker, Nicole Folstad, Matthew Ott, and Ratan Chopra for their assistance with experiments and for feedback on the manuscript. This material is based upon work supported by the National Science Foundation Graduate Research Fellowship under Grant No. 00006595 to KMD. Any opinions, findings, and conclusions or recommendations expressed in this material are those of the authors and do not necessarily reflect the views of the National Science Foundation. This project is funded by the USDA National Institute of Food Agriculture-Institute of Bioenergy, Climate and Environment, competitive grant no. 2014-67009-22305.

## Author Contributions

KMD and MDM conceived and planned the study. EBJ, ED, and MDM constructed the F2 mapping population. EBJ and ED identified the FLC-α allele. KMD designed and performed PCR experiments, whole genome sequencing and analyses, and comparative analyses. DLW collected all eMN designated pennycress accessions used in this study. KMD and MDM wrote the manuscript with feedback from all the co-authors.

## Supporting Information Brief Legends

**Fig. S1-** PCR strategy for identifying *flc-B* allele

**Fig. S2-** PCR strategy for identifying *flc-C* allele

**Fig. S3-** Peptide alignment of Brassicaceae FRIGIDA predicted peptides and pennycress FRI gene model

**Fig. S4-** Peptide alignment of Brassicaceae FLOWERING LOCUS C predicted peptides

**Fig. S5-** Genomic DNA alignment of pennycress FLOWER LOCUS C wild type and *flca* allele

**Fig. S6-** Identification and characterization of pennycress flc-α

**Table S1-** Summary of whole genome sequencing reads of pennycress accessions used in this study

**Table S2-** Primer sequences used in PCR experiments used in this study

**Table S3-** Summary of pennycress accessions and the corresponding alleles of FLC as identified in this study

